# Characterisation of gut microbiota of farmed Chinook salmon using metabarcoding

**DOI:** 10.1101/288761

**Authors:** Milica Ciric, David Waite, Jenny Draper, John Brian Jones

**Affiliations:** Ministry for Primary Industries, Investigation and Diagnostic Centre, Animal Health Laboratory, 66 Ward Street, Wallaceville, Upper Hutt, New Zealand; Ministry for Primary Industries, Investigation and Diagnostic Centre, Plant Health and Environment Laboratory, PO Box 2095, Auckland, New Zealand

## Abstract

With the growing importance of aquaculture worldwide, characterisation of the microbial flora of high-value aquaculture species and identification of gut flora shifts induced by changes in fish physiology or nutrition is of special interest.

Here we report the first metabarcoding survey of the intestinal bacteria of Chinook salmon (*Oncorhynchus tshawytscha*), an economically important aquacultured species. The microbiota of 30 farmed Chinook salmon from a single cohort was surveyed using metabarcode profiling of the V3-V4 hypervariable region of the bacterial 16S rRNA gene. Seawater, feed and intestinal samples, and controls were sequenced in quadruplicate to assess both biological and technical variation in the microbial profiles.

Over 1,000 operational taxonomic units (OTUs) were identified within the cohort, providing a first glimpse into the gut microbiota of farmed Chinook salmon. The taxonomic distribution of the salmon microbiota was reasonably stable, with around two thirds of individuals dominated by members of the family *Vibrionaceae*.

This survey was performed amid a summer heat wave, during which the fish exhibited reduced feeding. Although the sampled fish appeared healthy, they had minimal intestinal content, and the observed intestinal flora may represent the microbiota of fasting and stressed fish. Limited comparison between *Mycoplasma* and *Vibrio* sequences from the Chinook salmon gut and published microbial sequences from the intestines of a variety of fish species (including Atlantic salmon) indicated that despite the starvation and temperature variations, the replacement of *Vibrio* with *Mycoplasma* is occurring within expected ecological parameters and does not necessarily reflect colonisation by atypical microbes.

**DATA SUMMARY:** Raw sequences from Chinook salmon intestinal microbiome 16S survey generated on the Illumina platform are publically available through NCBI Sequence Read Archive (SRA) database:

Bioproject PRJNA421844

SRA study SRP134829

https://www.ncbi.nlm.nih.gov/bioproject/PRJNA421844

**IMPACT STATEMENT:** Although 16S metabarcoding surveys are becoming routine, little is still known about the microbiota of fish. This is the first survey of the intestinal microbiota of Chinook salmon, a species native to the Pacific Northwest which is farmed in New Zealand and Chile. While most intestinal microbiota studies are performed on faecal material, we directly sampled the intestine epithelium and content.

During the time of sampling, the farmed fish population was experiencing stress from a summer heat wave and had little intestinal content. Over 1,000 operational taxonomic units (OTUs) were identified within the intestines of the cohort, providing a first glimpse into the gut microbiota of farmed Chinook salmon.

We believe this survey will be of interest not only to those interested in fish biology and aquaculture, but also as an addition to the ongoing debate in the literature on sampling and DNA extraction methods for challenging samples.

## INTRODUCTION

The digestive tracts of all vertebrates harbour complex assemblages of microorganisms (microbial communities), collectively referred to as gut microbiota. The gut microbiota is an area of research interest universally applicable to Animalia, but the majority of studies on gut microbiota composition and function in vertebrates have been conducted in mammals. Comparatively little is known about the fish gut microbiota and its response to changing environmental conditions (1), despite the fact that fish represent roughly half of all living vertebrate species (2) and are of global economic significance.

The microbial community harboured in fish guts influences host physiology and is therefore of relevance to aquaculture. Gnotobiotic and conventional studies indicate the involvement of the gut microbiota in fish nutrition, development of intestinal epithelium, immunity, and disease (3). During the past few decades, substantial research has been carried out to characterize the gut microbiota in a wide range of fish species, focusing primarily on model organisms (i.e. zebrafish) and species relevant to aquaculture. As studies of fish gut microbial diversity have moved away from culture- and microscopy-based observations to the culture-independent molecular techniques, it has become clear that the intestinal microbiota of fish is more variable than previously realised. High throughput 16S rRNA sequencing has been increasingly employed to investigate changes in the fish intestinal microbiome caused by diet (including probiotics), starvation, pathogens, different lifestyles, and water temperature (1,4,5).

The fish gut harbours a combination of resident (autochthonous) microbiota, attached to the intestinal mucosa, and non-resident (allochthonous) microbiota comprised of microbes appearing transiently and/or associated with digesta (6,7). The composition of fish gut microbiota and species richness varies with life stage, diet, and environment (3,8) and differs between marine and freshwater species (9). Fish gut microbiota also varies between individuals, across the length of the gastrointestinal tract, and between gut content and mucosal surfaces (1,5,10–12). Interaction between time of sampling and diet is strongly related to the observed community structure (13). Nonetheless, phylogenetic and statistical analyses of 16S rRNA libraries suggest that core teleost gut communities may be shared between broad ranges of fish species (5,14,15).

Most fish gut microbiomes investigated to date comprise microbes from the phyla Actinobacteria, Bacteroidetes, Firmicutes, Fusobacteria, Planctomycetes, Proteobacteria and Tenericutes (1,3,16). The genera *Aeromonas* and *Pseudomonas* (Proteobacteria) and phylum Bacteroidetes dominate freshwater fish bacterial communities, while the genera *Vibrio*, *Pseudomonas* and *Alteromonas* (Proteobacteria) are reported to predominate the gut of marine fish (1,3,9). Proteobacteria and Firmicutes are the most reported phyla in the salmonid gut microbiome (7), although current knowledge of the bacterial diversity in the salmon gut is largely based on classical culturing techniques. Dominant culturable bacteria isolated from intestines of salmonid fish species include *Vibrio*, *Aliivibrio*, *Photobacterium*, *Lactobacillus*, *Lactococcus*, *Flavobacterium*, *Pseudomonas* and assorted *Enterobacteriaceae*. Recent sequencing-based studies have focused on salmonid species most prevalent in aquaculture - especially Atlantic salmon (*Salmo salar*) (6,17–22), Coho salmon (*Oncorhynchus kisutch*) (23), and rainbow trout (*Oncorhynchus mykiss*) (11,24–28). To our knowledge, no such studies have been published on Chinook salmon (*Oncorhynchus tshawytscha*), also known as king salmon, a species that is farmed in commercial quantities mainly in New Zealand and Chile.

The primary purpose of this study was to identify the baseline gut microbiota of farmed Chinook salmon and assess degree of variation among individuals within the same cohort. For this reason, our experimental design included duplication of the library indexing (to assess intra-run variation) and sequencing (to assess run-to-run variation) steps. We anticipated that this would enable us to differentiate between the effects of sequencing variation, if present, and biological variation between fish gut microbiota.

## METHODS

### Fish management

Chinook salmon used in this experiment were obtained from a commercial NZ King Salmon (NZKS) farm (Ruakaka Bay Farm in Queen Charlotte Sounds, New Zealand) during a standard harvest operation. All sampled fish were female, approximately 22 months old, and belonged to a single cohort. Individuals were reared in sea pens (20 m × 20 m × 15 m) using standard farm management practice (29). The fish were fed a Quinnat Plus 2200 commercial diet (BioMar, Denmark), delivered to the sea pens via a mobile hopper twice a day using a satiation feeding approach, until the harvest. No antibiotics, probiotics, antifungals, antivirals o antiparasitics were used during the rearing of the salmon.

### Sample collection and processing

Samples were collected in January 2015 (mid-summer) at the salmon farm from 30 apparently healthy Chinook salmon. The fish were harvested at an average weight of 3.6 kg using standard NZKS operational practice (29) and processed on the barge immediately after slaughter. The fish were dissected, and the length and the appearance of the mid-intestine, measured from the last pyloric caecae to the start of the distal intestine, was recorded. For each fish, a 1 cm section of the mid-intestine, including its gut content, was collected from the first half of the mid-intestines (roughly 2 cm past the pyloric caeca) using sterile instruments and into sterile tubes containing 5 ml RNAlater solution (Ambion, USA). Several feed pellets from the spinner supplying the relevant pen and 5 ml of seawater from the surface of the pen were also collected into sterile tubes containing 5 ml RNAlater solution. A sterile tube containing 5 ml RNAlater solution was handled identically to the rest of the samples, including being carried to the sampling site (barge) and briefly exposed to the air (negative control). The samples were transported on ice packs to the laboratory and stored at 4°C for two weeks prior to DNA extraction.

For each sample, the gut tissue section was removed aseptically from the RNAlater and opened longitudinally to release digesta using sterile forceps and scalpel, and the appearance of the gut content was recorded. The opened gut section was “washed” in 3 ml of RNAlater suspension in which it was stored by vigorous vortexing at maximum speed for 30 sec to release and homogenize gut content. The washed gut tissue was then aseptically removed, and the remaining homogenized sample was split into two 1.5 ml aliquots. One aliquot was used directly for the DNA extraction. For the other aliquot, prior to DNA extraction, material gently scraped from the mucosal surface of removed gut tissue was added to the sample to increase the likelihood of collecting bacterial cells adherent to gut epithelium or trapped in the mucus layer. Approximately 100-200 mg of feed pellets were crushed aseptically using mortar and pestle and homogenised in 1.5 ml of RNAlater by vortexing. The seawater sample was not processed prior to DNA extraction.

### DNA extractions

DNA was extracted from 1.5 ml aliquots of the salmon intestinal sample/RNAlater suspensions (one with and one without addition of scraped mucosal material), as well as from 1.5 ml of seawater/RNAlater suspension and 1.5 ml of feed/RNAlater suspension. Because of several possible sources of bacterial DNA contamination during sampling and DNA extraction (30), a DNA extraction was also performed from 1.5 ml of RNAlater carried during the sampling trip (negative control, to account for bacterial DNA being introduced from the DNA extraction and sample handling). DNA was extracted with NucleoSpin Soil kit (Macherey-Nagel, Germany) using an adapted manufacturer’s protocol and DNA yield, purity and integrity, as well as the presence of bacterial and host DNA in intestinal samples and controls were assessed as described in Supplementary methods (sub-heading ‘DNA extractions’).

### 16S rRNA amplicon library preparation and sequencing

DNA extracted from salmon intestinal samples with and without addition of scraped mucosal material were pooled for the amplification of the V3-V4 region of 16S rRNA gene. A series of template dilutions (1−, 2−, 5− and 10-fold dilutions) were tested for each sample and the dilution that produced the strongest 16S amplicon band, judged by agarose gel electrophoresis, was chosen for sequencing. For seawater and feed samples, the RNAlater, the no-template control (molecular water), and genomic DNA from the mock microbial community HM-782D (31), 16S amplicons were produced without template dilution. Detailed protocol on the preparation of sequencing library can be found in Supplementary methods (sub-heading ‘16S rRNA amplicon library preparation’).

Samples were normalised where possible to the equivalent concentration of 10 ng/µl and sent to the sequencing provider New Zealand Genomics Limited (NZGL) for indexing and sequencing. Indexing was performed by NZGL using the Nextera XT Index Kit (Illumina, USA) in duplicate for each sample, with two distinct dual index pairs used for each sample to assess the effect of index and intra-run variability. This dual indexing of the initial library of 35 samples (30 fish samples, seawater and feed samples, and 3 controls) resulted in a final sequencing library of 70 samples. All samples were pooled without normalization and sequenced twice using the Illumina MiSeq system with v3 reagents to produce 2 × 300 bp reads. Two separate 600-cycle 65-hour runs were performed using two MiSeq instruments to determine ‘run to run’ variation (technical replicates).

The reads were processed and assigned to OTUs using the QIIME software package, version 1.8.0 (32), as described in Supplementary methods (sub-heading ‘Sequence analysis’).

## RESULTS AND DISCUSSION

### 16S rRNA amplicon library

Intestinal samples ranged in length from 9 to 15 cm and had mostly normal appearance. Although the sampled salmon had been fed as per usual farm protocol until the harvest, only 8 contained visible feed content. Sampling was performed during a mid-summer water temperature spike, and salmon tend to go off feed when water temperatures are high (33). Following completion of this study, salmon farms in the region, including the one surveyed, reported elevated fish mortality (>30%) over the sampling period, that was likely associated with this stressor.

Quality checks of the DNA samples extracted from salmon intestinal samples, seawater, feed and RNAlater (control), as well as of the 16S rRNA amplicons are discussed in Supplementary results (sub-heading ‘16S rRNA amplicon library QC’).

Although high levels of host DNA complicated QC and sample normalisation during sequencing library preparation, it was not anticipated this would have a major effect on the sequencing run itself, as the contaminating DNA lacks the sequencing adapters. Amplicons from 16S V3-V4 sequencing library, produced from the 30 salmon intestinal samples, seawater and feed samples and controls (RNAlater, no-template control and mock microbial community), were indexed and sequenced on a MiSeq instrument by NZGL without further processing or normalisation, except diluting higher-concentration samples to 10 ng/µl. As a result, the sequencing library contained uneven concentrations of amplicons across the samples, and we attempted to assess the effect of this difference on the resulting microbiota profiles.

### Sequencing metrics and analysis

Sequencing yielded a total of 18 million 16S rRNA gene amplicon sequences, combined over both sequencing runs. Approximately 10.2 million and 7.9 million of reads were assigned to an index in the first and the second sequencing run. Following read pair joining, removal of short sequences and stringent quality filtering, sequence numbers were reduced to 750 thousand sequences spread over 140 samples (Table 1). A full list of all OTUs identified during quality control stages is reported in Supplementary tables (Table S1). Overall sequencing error rate, based on sequencing mock microbial community, was 0.49%. The number of sequences recovered, and OTU richness in all the controls (RNAlater, no-template control and mock microbial community) was low indicating a paucity of contaminating DNA sequences in these controls.

**Table 1.**
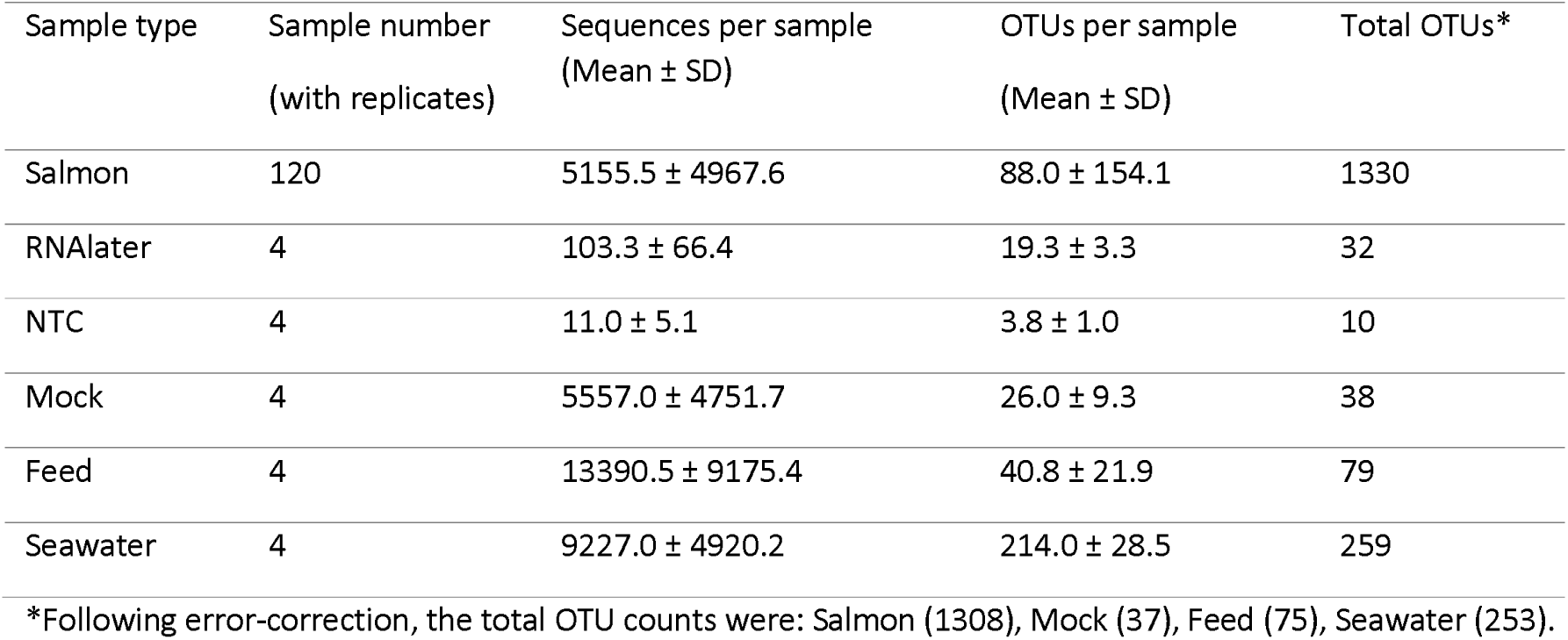
Summary statistics for the sequencing run.

| Sample type | Sample number<br>(with replicates) | Sequences per sample<br>(Mean ± SD) | OTUs per sample<br>(Mean ± SD) | Total OTUs* |
| --- | --- | --- | --- | --- |
| Salmon | 120 | 5155.5 ± 4967.6 | 88.0 ± 154.1 | 1330 |
| RNAlater | 4 | 103.3 ± 66.4 | 19.3 ± 3.3 | 32 |
| NTC | 4 | 11.0 ± 5.1 | 3.8 ± 1.0 | 10 |
| Mock | 4 | 5557.0 ± 4751.7 | 26.0 ± 9.3 | 38 |
| Feed | 4 | 13390.5 ± 9175.4 | 40.8 ± 21.9 | 79 |
| Seawater | 4 | 9227.0 ± 4920.2 | 214.0 ± 28.5 | 259 |

\*Following error-correction, the total OTU counts were: Salmon (1308), Mock (37), Feed (75), Seawater (253).

Results for analysis of variation are available in Supplementary results (sub-heading ‘Analysis of variation’).

### Microbiota of farmed Chinook salmon

A total of 1,308 OTUs, each corresponding roughly to a bacterial species, were identified within the intestinal microbiota of the 30 salmon. The salmon microbiota was typically dominated by several abundant bacterial lineages, which skewed the evenness of the community (Fig. 1; Supplementary tables (Table S2)).

**Fig. 1.**
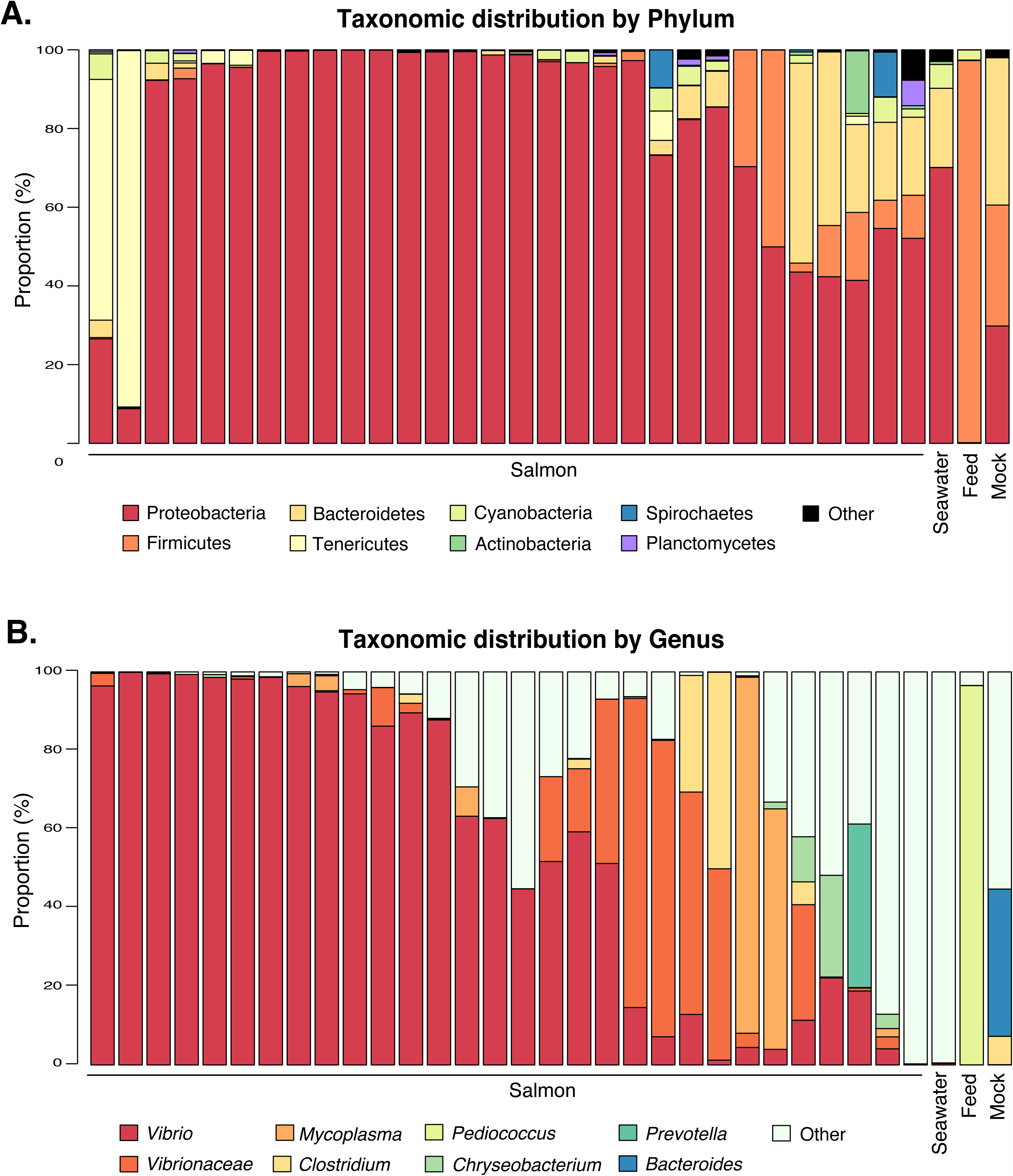
Phylum (A) and genus (B) level taxonomic distribution of microbiomes surveyed. Bars report the mean abundance for each individual sample. The top 9 most abundant genera (across all samples) are reported, all others are aggregated into ‘Other’.

The taxonomic distribution of the salmon microbiota was conserved between individuals, but notably different between salmon and environmental samples (Fig. 1). The intestinal microbiota of the majority of fish was dominated by the family *Vibrionaceae*, as has been observed in Atlantic salmon (Salmo salar) (22). Interestingly, two of the thirty sampled individuals had a microbiota dominated by *Mycoplasma*; dominance of *Mycoplasma* spp. has been seen before in wild-caught Atlantic salmon (18). Phylogenetic inference of OTUs belonging to these lineages demonstrated that these OTUs are closely related to species identified in similar asnalyses of fish microbiomes (14,18,34–37) (Fig. S1 and Fig. S2). This finding indicates that despite the starvation and temperature variations, the observation of *Vibrio* and *Mycoplasma* is consistent with typical fish-associated species and are unlikely to represent colonisation by novel lineages. Individual variation in salmon gut microbiota is further discussed in Supplementary results (sub-heading ‘Diversity and individual variation in salmon gut microbiota’).

When attempting to characterise a “core microbiome” of the sampled salmon, it was apparent that at the level of individual OTUs, the microbiota of the fasting fish is quite variable within this single cohort. This is not surprising, as conservation of intestinal microbiota occurs primarily at the level of metabolic function, while the specific bacterial species fulfilling that function within an individual animal can vary significantly (38,39). Only OTUs belonging to the family *Vibrionaceae* were present in over 90% of individuals sampled. The other OTUs present in at least 60% of individuals represented the genera *Synechococcus*, *Clostridium*, *Pseudomonas*, *Chryseobacterium*, *Brevundimonas*, *Sphingomonas*, *Paracoccus*, NS5 marine group, NS4 marine group, *Polaribacter*, *Acinetobacter*, *Sulfitobacter*, *Loktanella*, *Photobacterium*, and *Pseudoalteromonas*, or the families *Rhodobacteraceae*, *Rickettsiales* SAR116_clade, *Rhizobiales*, and *Oceanospirillales*, where a specific genus name could not be assigned.

A recent study of the effects of starvation on the Asian seabass (*Lates calcarifer*), another farmed carnivorous species, reported a major shift in the intestinal microbiota towards members of the phylum Bacteroidetes, driven by an increase in members of the classes *Sphingobacteria* and *Bacteroidia* and a decrease in the *Betaproteobacteria* (40). Without a control group of salmon feeding normally, it is impossible to determine if a similar effect is occurring in this data, but if so, this shift is not apparent in the core microbiota of the sampled cohort. Members of Bacteroidetes in the salmon only approached the levels of Proteobacteria in 10% of individuals (3 fish).

Although the microbiota of the pellet feed was dominated by a strain of *Pediococcus* (Fig. 1; median abundance 96.5%), this organism was almost undetectable in the salmon gut (<0.01%). *Pediococcus acidilactici* strain MA 18/5M is sold as the probiotic product Bactocell® (Lallemand Inc., Canada) for reducing intestinal inflammation in fish (41), and this product is added to some BioMar feed pellets (BioMar Group, Denmark). The low abundance of this organism in the intestine suggests that the organism is not able to establish in the gut and quickly declines when salmon are not actively consuming the probiotic.

Importantly to aquaculture of these fish, an OTU matching *Piscirickettsia salmonis* was detected in four individuals. This is consistent with the recent finding of *P. salmonis* - like bacteria at multiple salmon farms in the Marlborough Sounds (42,43), including the farm sampled during this study. Accurate taxonomic identification from partially sequenced 16S rRNA gene fragments is problematic (44,45), but as *P. salmonis* is a known salmon pathogen (46), this detection demonstrates the applicability of metabarcoding for detection of potential pathogens in asymptomatic individuals and highlights the need to follow up suspicious findings with targeted diagnostic tests.

In conclusion, we conducted the first metabarcoding survey of the bacterial intestinal microbiota of farmed Chinook salmon. Over thousand OTUs were identified within the intestines of a cohort of 30 fish, providing a first glimpse into the gut flora of this aquacultured species. Our survey was performed during a summer water temperature spike, during which the salmon had stopped eating, and salmon farms in the region (including the one surveyed) were experiencing mortality (>30%) associated with this stressor. As a result, although the sampled fish appeared healthy, their intestinal flora likely represents the microbiota of fasting and stressed fish. It is unknown whether the microbial profiles reported here would remain similar for fish without this environmental stressor.

## Acknowledgements

The authors would like to thank Dr Colin Johnston from Aquaculture New Zealand and Mark Preece from New Zealand King Salmon for providing salmon for sampling; as well as Dr Suzanne Keeling, Dr Lia Liefting and Andrew Bell from the MPI for useful suggestions during writing.

The following reagent was obtained through BEI Resources, NIAID, NIH as part of the Human Microbiome Project: Genomic DNA from Microbial Mock Community B (Even, Low Concentration), v5.1L, for 16S rRNA Gene Sequencing, HM-782D.

## Ethical statement

This work did not require ethical approval.

The fish examined in this research were obtained from the commercial salmon farm (NZ King Salmon farm) after normal commercial harvest and prior to gutting and gilling. All fish were harvested using standard NZ King Salmon operational practice (29). Under New Zealand’s Animal Welfare Act 1999, dissections on carcass material do not require approval by the animal ethics committee.

## ABBREVIATIONS

NMDS: non-metric multidimensional scaling
OTU: operational taxonomic unit
PERMANOVA: permutational multivariate analysis of variance
QC: quality control

