## Supplementary Materials for "Characterisation of gut microbiota of farmed Chinook salmon using metabarcoding"

**SUPPLEMENTARY TEXT**

**SUPPLEMENTARY METHODS**

**DNA extractions**

Each sample was vortexed briefly before transfer into NucleoSpin bead tubes using a wide-bore pipette tip (the top of the tip was cut off using sterile scissors). Due to the limitation in volume that can be accommodated by components of the kit, for each sample three 0.5 ml sample loads were processed sequentially through a single bead beating tube, inhibitor removal column and DNA binding column, for a total of 1.5 ml processed sample per extraction. For each 0.5 ml of sample loaded, we used the following sequence of processing steps. Samples were mixed with 0.7 ml of lysis buffer SL2 (without addition of enhancer) and subjected to horizontal bead-beating for 10 mins at room temperature at maximum speed using the Vortex-Genie 2 adaptor (MO BIO Laboratories, USA). After spinning down beads and cell debris, the supernatant was transferred to a sterile 2 ml Eppendorf tube and incubated on ice for 15 mins with 180 µl of precipitation buffer SL3. After centrifugation, the supernatant was put through the NucleoSpin Inhibitor Removal column. The filtrate was transferred to a new 2 ml tube and 500 µl of binding buffer SB was added prior to loading on a DNA binding column. After sequential binding of DNA from all three 0.5 ml aliquots of the starting 1.5 ml sample to the column, washing steps were performed according to manufacturer’s protocol. To increase DNA yield, DNA was eluted sequentially with 2 X 30 µl of elution buffer SE that was pre-heated at 80°C.

The DNA purity was assessed by determining the absorbance ratio of nucleic acids and other contaminating molecules (A260/A280 and A280/A230) using the NanoDrop spectrophotometer 8000 (Thermo Fisher Scientific, USA). The DNA yield was measured using the Qubit fluorometer (Thermo Fisher Scientific, USA). The DNA integrity was assessed by gel electrophoresis on a 1% agarose gel in comparison to high molecular weight DNA standards (40 kb Fosmid Control DNA (Epicentre, USA) or 48.5 kb Lambda DNA (Thermo Fisher Scientific, USA)).

The presence of bacterial and host DNA in intestinal samples and controls was assessed by singleplex 16S and 18S real-time PCR, respectively. DNA extracted from the intestinal samples were normalised to 30 ng (2 µl) and used as template for PCR reaction. DNA extracted from RNAlater, seawater and feed samples were not normalised and 2 µl of DNA sample were used for each PCR reaction. The 16S and 18S real-time PCR were performed simultaneously on the same plate using PerfeCTa qPCR ToughMix (Quantabio, USA), containing 50nM of each forward and reverse primers and fluorogenic probe for either 16S (1) or 18S (2) targets in a final volume of 10 µL. PCR conditions were as follows: initial denaturation at 95°C for 2 min, followed by 40 cycles of denaturing (95°C for 30 sec) and annealing (60°C for 30 sec). All PCR runs included a positive (*E. coli* ATCC25922 genomic DNA for 16S and human cell line DNA for 18S) and negative (no template) controls.

**16S rRNA amplicon library preparation**

The first step of the 16S rRNA amplicon library preparation was performed using the primer pair targeting V3-V4 region of the 16S rRNA gene (3). A touchdown protocol and preheated thermal cycler were used to minimize unspecific primer binding. PCR amplification was performed in duplicate 25 µl reactions in two separate runs (4 reactions per sample in total), using 2.5 µl template DNA at the optimum dilution for each sample. The total template DNA input in PCR reactions ranged from 19.3 ng-462.5 ng. Reactions also contained a final concentration of 200 µM dNTPs, 0.2 µM primers and 0.02 U/µl of Phusion Hot Start II High-Fidelity DNA Polymerase (Life Technologies, USA). A total of 30 cycles of PCR amplification were performed as follows: initial denaturation at 98°C for 3 min, 15 touchdown cycles of denaturing (98°C for 30 sec), annealing (62-55°C for 30 sec, Δt 0.5°C) and elongation (72°C for 40 sec), followed by 15 cycles of denaturing (98°C for 30 sec), annealing (55°C for 30 sec) and elongation (72°C for 40 sec), and then a final 5 min extension at 72°C.

The 4 PCR reactions for each sample were pooled, and 90 µl was cleaned using Agencourt AMPure XP (Beckman Coulter, USA) beads according to the MiSeq protocol (4) and eluted with 40 µl 10 mM Tris (pH 8.5). Amplicon size and effectiveness of the bead clean-up was analysed by agarose gel electrophoresis. Densitometry analysis of gel images, using the software package ImageJ (5), was performed to quantify amplicon band intensities relative to the *E.coli* control and estimate amplicon yield. The amount of total DNA in each sample, representing a mixture of amplicons and template DNA, was quantified fluorometrically by Qubit.

**Sequence analysis**

Initial sequence analysis was performed using the QIIME software package, version 1.8.0 (6). Adapter and barcode sequences were removed from raw amplicon sequences using cutadapt version 1.1.0 (7) and paired end reads were then joined using the QIIME fastq-join method, simultaneously filtering sequences for Q ≥30. Sequencing error rates were quantified using the seq.error command in mothur (8). Briefly, sequences from the mock community were extracted from the full data set and aligned to the SILVA SSU database (9,10), release 119 (11). A reference database of 16S rRNA sequences from the mock community (Microbial Mock Community B, BEI Resources) were obtained and aligned using the same process. Reference sequences were then trimmed to the same alignment space as the amplicons, and sequence error between reference and amplicons calculated. Sequences were clustered into operational taxonomic units (OTUs) of 97% sequence similarity using the *de novo* method and screened for chimeras using USEARCH algorithm (12,13). A representative sequence for each OTU was taxonomically classified against the SILVA SSU database release 119 (11) using the assign_taxonomy.py command. Taxonomy was assigned as the consensus taxonomy of the top 10 hits per sequence and any sequences identified as mitochondria or archaea were removed from the dataset, as were sequences that could not be classified to the domain level. The remaining sequences were aligned to the 16S ribosomal RNA secondary structure using PyNAST (14) and a phylogenetic tree was built using FastTree (15).

In order to limit the effects of accidental contamination during sample handling, data were analysed in two ways. First, the complete data set was analysed without accounting for bacterial sequences identified in the no template control samples (‘uncorrected data’). Second, all OTUs observed in the no template control and RNAlater blank extraction samples, as well as OTUs identified as likely artefacts due to their inverse correlation with amplicon concentration (16) were removed from the data set (‘error-corrected data’).

Taxonomic summaries were generated using QIIME, and alpha diversity was calculated using the Chao1 richness estimator and the Shannon and inverse Simpson diversity estimators. Alpha diversity calculations were performed using the error-corrected data set with sampling depth rarefied to 500 and 1,000 sequences per sample, as well as without subsampling (17). Beta-diversity (between-sample) calculations were performed in mothur (8). In order to minimise the impact of artefacts resulting from random subsampling, each sample was randomly subsampled 1,000 times and beta diversity measures calculated from each iteration. The average distance was then reported for each metric. Beta-diversity was calculated using presence/absence (Jaccard and unweighted UniFrac distance) and relative abundance (Bray-Curtis, Yue-Clayton theta and weighted UniFrac distance) measures for each data combination and subsampling depth. To ascertain the effect of technical biases on the salmon microbiota, data were analysed using permutational multivariate analysis of variance (PERMANOVA) to quantify the contribution of different metadata factors to the beta-diversity of the samples (18,19). PERMANOVA was performed in the R software environment using the vegan package (20), and data were visualised using non-metric multidimensional scaling (NMDS) (21,22). The effects of microbiome donor, sequencing run, index, amplicon yield (determined by both Qubit and densitometry) and sequences obtained were tested against each beta-diversity matrix with 10,000 permutations to determine the statistical significance of each variable on the microbiome structure.

OTUs obtained from the Chinook salmon microbiota were compared to those obtained from previously published research studying the microbiota of various fish species (23–29). Where 16S rRNA gene amplicon data were provided by authors, raw sequence reads were obtained from the NCBI Sequence Read Archive and OTUs generated using the protocol detailed above. Sequences classified as belonging to bacterial lineages of interest were aligned using the SINA web aligner (30) and inserted into the SILVA database (10) (version 128, non-redundant). All alignments were manually inspected and corrected for errors. Representative sequences belonging to OTUs, clones, and cultivated type material were selected and masked using the SILVA ‘ssuref:bacteria’ filter. Phylogenetic inference was then performed using a maximum likelihood approach on sequences of ≥1,000 bp length using IQ-Tree (31), under the general time reversible model with gamma-distributed rate heterogeneity (GTR+G) and 1,000 bootstrap re-samplings to assess node stability. Short length sequences (<1,000 bp length) were inserted into the tree using the ‘ARB Parsimony’ tool in the ARB software environment (32).

**SUPPLEMENTARY RESULTS**

**16S rRNA amplicon library QC**

A quality check of the DNA extracted from salmon intestinal samples, seawater, feed and RNAlater (control) showed A260/A280 ratios in the range of 1.78 - 1.94 and A260/A230 ratios in the range of 1.1 - 2.2. The DNA samples extracted from the salmon gut content were dominated by a high molecular weight DNA, whereas DNA extracted from the seawater, feed samples and RNAlater (control) were of low concentration and were not visible on an agarose gel. Using 16S/18S real-time PCRs, a highly variable and often low quantity of bacterial DNA relative to host DNA was observed, which is consistent with the sparse intestinal content in the majority of samples. For 16S Cq values ranged from 20.3 to 36.7, and for 18S Cq values ranged from 14.2 to 24.6.

To address vastly different amplicon yields for individual samples, the final PCR to generate the 16S rRNA amplicon library used an optimised template concentration for each sample, as described in methods section (‘16S rRNA amplicon library preparation and sequencing’ ). This was problematic for performing standard quality control (QC) measurements on amplicon samples that were subsequently used for the 16S rRNA amplicon library preparation. Amplicon yield could not be reliably measured fluorometrically or spectrophotometrically for all samples due to high template carryover, given that the initial concentration of template DNA in the PCR reactions was up to 18.5 ng/µl. The Bioanalyzer instrument (Agilent Technologies, USA) also could not be used to assess the library as per standard sequencing protocol due to the genomic/host DNA contamination. In the Bioanalyzer instrument, all of the samples are run sequentially through a single capillary; therefore the slowly-migrating genomic DNA bands overlap with the next sample running through the capillary, and it becomes impossible to match bands to samples. For these reasons, we had to rely primarily on gel electrophoresis for QC. To generate an estimate of relative amplicon concentration unaffected by host DNA concentration, a densitometry analysis was performed as described in Supplementary methods (sub-heading ‘16S rRNA amplicon library preparation’).

**Analysis of variation**

Five measures of community structure (Jaccard, unweighted UniFrac, Bray-Curtis, Yue-Clayton theta and weighted UniFrac distances) were computed from the final OTU table. Measures were performed using both the full (‘uncorrected’) and ‘error-corrected’ data table. Boxplot visualisation of the PERMANOVA data (distribution of fit measurements according to metadata characteristics for all calculated beta diversity measures, reported according to error correction strategy) showed high variability (Fig. S3). High variability is due to the aggregation of multiple beta diversity scores using multiple subsampling strategies. For example, unweighted UniFrac typically performed poorly in the individual category (median R2 = 0.42), while Yue-Clayton provided an excellent fit for data (median R2 = 0.97). Under all 5 distance metrics tested, the effect of the biological variables (individual variation, sample type) in the study were profoundly greater than the technical aspects - sequencing run, index, and amplicon yield. Although the individual donor and amplicon yield variables are highly confounded, manually identifying and analysing a subset of the data with highly similar amplicon yields (as determined by densitometry) showed the same pattern as for the full data set (data not shown). The data demonstrate that the microbiota donor is the strongest contributor towards community structure, and that technical parameters quantified within this study do not significantly influence patterns in community structure.

Visualisation of this data performed using a NMDS plot of community structure (Fig. S4), based on Yue-Clayton theta distance with no subsampling, showed that the 4 technical replicates for each sample clustered together, while feed and seawater samples and controls (NTC, RNAlater, mock) were distinct. With the exception of the RNAlater control in the ‘uncorrected data’, the microbiota present in the additional samples (feed, seawater) did not have a significant effect on the fish gut microbiota, although it is likely that these sources act to seed the gut microbiota with certain microbial lineages.

Overall, it appears that the effects of technical variation, including amplicon yield, indexing, and MiSeq run, when each step is performed by the same individual and instrument, are negligible. Contamination by reads from the kits and reagents, however, does appear to have a minor effect and should be accurately quantified and removed from the data prior to analysis.

A list of all OTUs identified during quality control stages is reported in Supplementary tables (Table S1). Removal of OTUs associated with negative controls or with distributions inversely correlated to amplicon yield (16) improved the fit of the data (Fig. S3).

Overall, although the impact of contamination is negligible on the level of gross variation between individuals, it does have an effect. Batch to batch variations of kit and reagent microbial contaminant profiles were previously reported (33,34); therefore we highly recommend running such controls for every individual kit and PCR reagent batch used.

**Diversity and individual variation in salmon gut microbiota**

The gut microbiota of sampled salmon was less diverse than seawater and also contained a lower species richness (Fig. S5). This finding is expected, as the acquisition and maintenance of fish gut microbiota is a complex process. It is driven by both environmental availability of potential microbial colonisers and host physiological pressures in a highly selective gut environment (35,36).

The presence of gut digesta at the time of sampling was the only parameter with observable effect and although presence of digesta did not drive a strong separation of salmon samples, there was a tendency for these samples to cluster within the broader salmon data (Fig. S6). Such a pattern is not unexpected, as the digesta is the primary source of nutrients for the gut microbiota, thus affecting the microbial community structure. The apparent lack of similarity between the microbiota structure of the fed salmon gut and the feed pellets provides evidence that the food source is not contributing significantly to the colonising gut microbiota.

**SUPPLEMENTARY FIGURES AND TABLES**

### Supplementary figures

**Fig. S1. Phylogenetic analysis of *Vibrio*-like 16S rRNA sequences obtained from marine vertebrates.** Sequences in black were obtained from cultivated *Vibrio* isolates, and those in grey from publicly available microbiome surveys. Sequences in red are *Vibrio*-like OTUs from this data set. Dashed lines represent short sequences (<1,000 bp) inserted into the fixed tree. Bootstrap support is represented by solid (≥90%) and hollow (≥75%) junctions. Scale bar represents 10% sequence divergence.

**Fig. S2. Phylogenetic analysis of *Mycoplasma*-like 16S rRNA sequences obtained from marine vertebrates.** Sequences in black were obtained from cultivated *Mycoplasma* isolates, and those in grey from publicly available microbiome surveys. Sequence in red represents the *Mycoplasma*-like OTU from this data set. Dashed lines represent short sequences (<1,000 bp) inserted into the fixed tree. Bootstrap support is represented by solid (≥90%) and hollow (≥75%) junctions. Scale bar represents 10% sequence divergence.

**Fig. S3. Average fit of data based on metadata characteristics.** Boxplot visualisation of the distribution of fit measurements according to metadata characteristics for all calculated beta diversity measures. Data are reported according to error correction (i.e. contaminant removal) strategy, as defined in section 2.4. High variability is due to the aggregation of multiple beta diversity scores using multiple subsampling strategy. For example, unweighted UniFrac typically performed poorly in the Individual category (median R2 = 0.42), while Yue-Clayton provided an excellent fit for data (median R2 = 0.97). Nonetheless, it is clear that the majority of variation is attributable to variation between individuals.

**Fig. S4. NMDS plots of community structure**. Plots are based on Yue-Clayton theta distance with no subsampling. Shadowing reflects clustering of individual samples. Left: Uncorrected data (stress = 0.24, r2 = 0.95). Right: Error-corrected data (stress = 0.23, r2 = 0.95). Controls (NTC, Mock, Feed, Sea water, RNAlater) cluster separately; the remaining clusters tend to be composed of the 4 technical replicates from an individual fish.

**Fig. S5. Comparison of within-sample diversity.** Comparison of alpha (within-sample) diversity parameters grouped by sample type, subsampled to 1,000 sequences per sample. Left: Chao1 richness estimator. Middle: Shannon diversity estimator. Right: Inverse Simpson diversity estimator.

**Fig. S6. NMDS plot of salmon gut microbiota with the presence of gut content recorded.** Data presents the error-corrected data set without subsampling, and samples are coloured based on sample type. Ellipses represent the 95% confidence interval around all samples with recorded gut content. Left: Jaccard distance (stress = 0.20, r2 = 0.96). Right: Yue-Clayton theta distance (stress = 0.23, r2 = 0.95).

### Supplementary tables

**Table S1. Contaminant OTUs identified in the 16S sequence data set of sampled Chinook salmon.**

| **OTUs identified in NTC and RNAlater samples** | | | | |
| --- | --- | --- | --- | --- |
| **Phylum** | **Class** | **Order** | **Family** | **Genus** |
| Firmicutes | *Bacilli* | *Lactobacillales* | *Carnobacteriaceae* | *Atopostipes* |
| Proteobacteria | *Betaproteobacteria* | *Burkholderiales* | *Burkholderiaceae* | *Burkholderia* |
|  |  |  | *Comamonadaceae* | *Schlegelella* |
|  |  |  |  | *Tepidimonas* |
|  | *Gammaproteobacteria* | *Oceanospirillales* | SAR86_clade |  |
|  |  | *Pseudomonadales* | *Pseudomonadaceae* | *Pseudomonas* |
|  |  | *Vibrionales* | *Vibrionaceae* | *Vibrio* |
|  | *Epsilonproteobacteria* | *Campylobacterales* | *Helicobacteraceae* | *Helicobacter* |
| **OTUs identified in Jervis-Bardy corrections** | | | | |
| **Phylum** | **Class** | **Order** | **Family** | **Genus** |
| Firmicutes | *Bacilli* | *Bacillales* | *Bacillaceae* | *Bacillus* |
|  |  |  | *Staphylococcaceae* | *Staphylococcus* |
|  |  | *Lactobacillales* | *Carnobacteriaceae* | *Atopostipes* |
| Proteobacteria | *Alphaproteobacteria* | *Rhizobiales* | *Bradyrhizobiaceae* | *Bosea* |
|  |  |  |  | *Bradyrhizobium* |
|  |  |  | *Hyphomicrobiaceae* | *Hyphomicrobium* |
|  |  |  | *Methylobacteriaceae* | *Methylobacterium* |
|  |  |  | *Phyllobacteriaceae* | *Mesorhizobium* |
|  |  | *Sphingomonadales* | *Sphingomonadaceae* | *Novosphingobium* |
|  |  |  |  | *Sphingomonas* |
|  |  |  |  | *Sphingopyxis* |
|  | *Betaproteobacteria* | *Burkholderiales* | *Burkholderiaceae* | *Burkholderia* |
|  |  |  |  | *Cupriavidus* |
|  |  |  | *Comamonadaceae* | *Curvibacter* |
|  |  |  |  | *Schlegelella* |
|  | *Gammaproteobacteria* | *Oceanospirillales* | *Halomonadaceae* | *Halomonas* |
|  |  | *Pseudomonadales* | *Moraxellaceae* | *Acinetobacter* |
|  |  |  |  | *Enhydrobacter* |
|  |  |  | *Pseudomonadaceae* | *Pseudomonas* |
|  |  | *Xanthomonadales* | *Xanthomonadaceae* | *Stenotrophomonas* |

Note: Because an OTU roughly corresponds to a single species (based on sequence identity), many distinct OTUs can belong to the same phylogenetic groups at the genus level or above. For example, the dominant *Vibrio* OTUs observed in the fish and seawater samples were distinct from *Vibrio* OTUs observed in the controls.

**Table S2. The most frequent bacterial taxa found in the intestine of the sampled Chinook salmon.**

| **Phylum** | **Class** | **Order** | **Family** | **Average**  **abundance** | | **Coverage** |
| --- | --- | --- | --- | --- | --- | --- |
| Proteobacteria |  |  |  | 83.1 | | 100 |
|  | *Alphaproteobacteria* |  |  | 5.4 | | 97 |
|  |  | *Caulobacterales* |  | 0.1 | | 87 |
|  |  |  | *Caulobacteraceae* | 0.1 | | 77 |
|  |  | *Rhizobiales* |  | 0.6 | | 83 |
|  |  |  | *Hyphomicrobiaceae* | 0.1 | | 53 |
|  |  |  | *Phyllobacteriaceae* | 0.2 | | 67 |
|  |  | *Rhodobacterales* | *Rhodobacteraceae* | 3.9 | | 93 |
|  |  | *Rhodospirillales* |  | 0.1 | | 77 |
|  |  |  | *Rhodospirillaceae* | 0.1 | | 70 |
|  |  | *Rickettsiales* |  | 0.1 | | 77 |
|  |  |  | *SAR116_clade* | 0.0 | | 53 |
|  |  | *Sphingomonadales* | *Sphingomonadaceae* | 0.2 | | 67 |
|  | *Betaproteobacteria* |  |  | 0.4 | | 93 |
|  |  | *Burkholderiales* |  | 0.2 | | 90 |
|  |  |  | *Oxalobacteraceae* | 0.0 | | 57 |
|  | *Deltaproteobacteria* |  |  | 0.2 | | 57 |
|  | *Epsilonproteobacteria* |  |  | 0.0 | | 37 |
|  | *Gammaproteobacteria* |  |  | 77.0 | | 100 |
|  |  | *Alteromonadales* |  | 1.2 | | 87 |
|  |  |  | *Alteromonadaceae* | | 0.8 | 80 |
|  |  |  | *Pseudoalteromonadaceae* | | 0.4 | 73 |
|  |  | *Enterobacteriales* | *Enterobacteriaceae* | | 0.0 | 53 |
|  |  | *Oceanospirillales* |  | 0.4 | | 83 |
|  |  |  | *SAR86_clade* | 0.1 | | 73 |
|  |  | *Pseudomonadales* |  | 0.6 | | 90 |
|  |  |  | *Moraxellaceae* | 0.5 | | 80 |
|  |  |  | *Pseudomonadaceae* | 0.1 | | 70 |
|  |  | *Thiotrichales* |  | 0.1 | | 50 |
|  |  | *Vibrionales* | *Vibrionaceae* | 74.0 | | 100 |
|  |  | *Xanthomonadales* |  | 0.3 | | 50 |
| Bacteroidetes |  |  |  | 3.0 | | 100 |
|  | *Bacteroidia* |  |  | 0.4 | | 47 |
|  | *Cytophagia* |  |  | 0.3 | | 43 |
|  | *Flavobacteriia* | *Flavobacteriales* | *Flavobacteriaceae* | 1.7 | | 100 |
|  | *Sphingobacteriia* | *Sphingobacteriales* | *Chitinophagaceae* | 0.0 | | 50 |
| Firmicutes |  |  |  | 4.3 | | 100 |
|  | *Bacilli* |  |  | 0.7 | | 87 |
|  |  | *Bacillales* | *Bacillaceae* | 0.0 | | 60 |
|  |  | *Lactobacillales* | *Streptococcaceae* | 0.1 | | 53 |
|  | *Clostridia* | *Clostridiales* | *Clostridiaceae_1* | 3.4 | | 77 |
| Cyanobacteria |  |  |  | 1.3 | | 97 |
|  | *Cyanobacteria* | *Chloroplast* |  | 0.7 | | 93 |
|  |  | *Cyanobacteria Subsection I* | | 0.4 | | 80 |
|  | *Cyanobacteria* |  |  | 0.6 | | 87 |
| Actinobacteria |  |  |  | 0.2 | | 80 |
|  | *Acidimicrobiia* | *Acidimicrobiales* |  | 0.1 | | 50 |
|  | *Actinobacteria* |  |  | 0.1 | | 67 |
|  |  | *Micrococcales* |  | 0.1 | | 53 |
| Verrucomicrobia |  |  |  | 0.2 | | 67 |
|  | *Verrucomicrobiae* | *Verrucomicrobiales* | *Verrucomicrobiaceae* | 0.0 | | 57 |
|  | *Opitutae* |  |  | 0.0 | | 37 |
| Tenericutes | *Mollicutes* | *Mycoplasmatales* | *Mycoplasmataceae* | 6.6 | | 63 |
| Planctomycetes |  |  |  | 0.6 | | 60 |
| Candidate division OM190 | | | | 0.0 | | 27 |
|  | *Planctomycetacia* | *Planctomycetales* | *Planctomycetaceae* | 0.5 | | 60 |
| Spirochaetes | *Spirochaetes* |  |  | 0.2 | | 40 |
| Chlamydiae | *Chlamydiae* |  |  | 0.0 | | 37 |
| Candidate division TM7 | | | | 0.2 | | 33 |
| Acidobacteria | *Holophagae* |  |  | 0.0 | | 27 |
| Candidate division BD1-5 | | | | 0.1 | | 27 |
| Candidate division TM6 | | | | 0.0 | | 27 |

Average abundance: the average percent of reads per sample matching that taxon; Coverage: the percent of individuals carrying organisms from that taxon. Coverage cut-offs for inclusion in this table are as follows: Phylum & Class >25%, Order > 50%, Family >50%.
