## Supplementary figures and images for "Characterisation of gut microbiota of farmed Chinook salmon using metabarcoding"

### Supplementary Materials

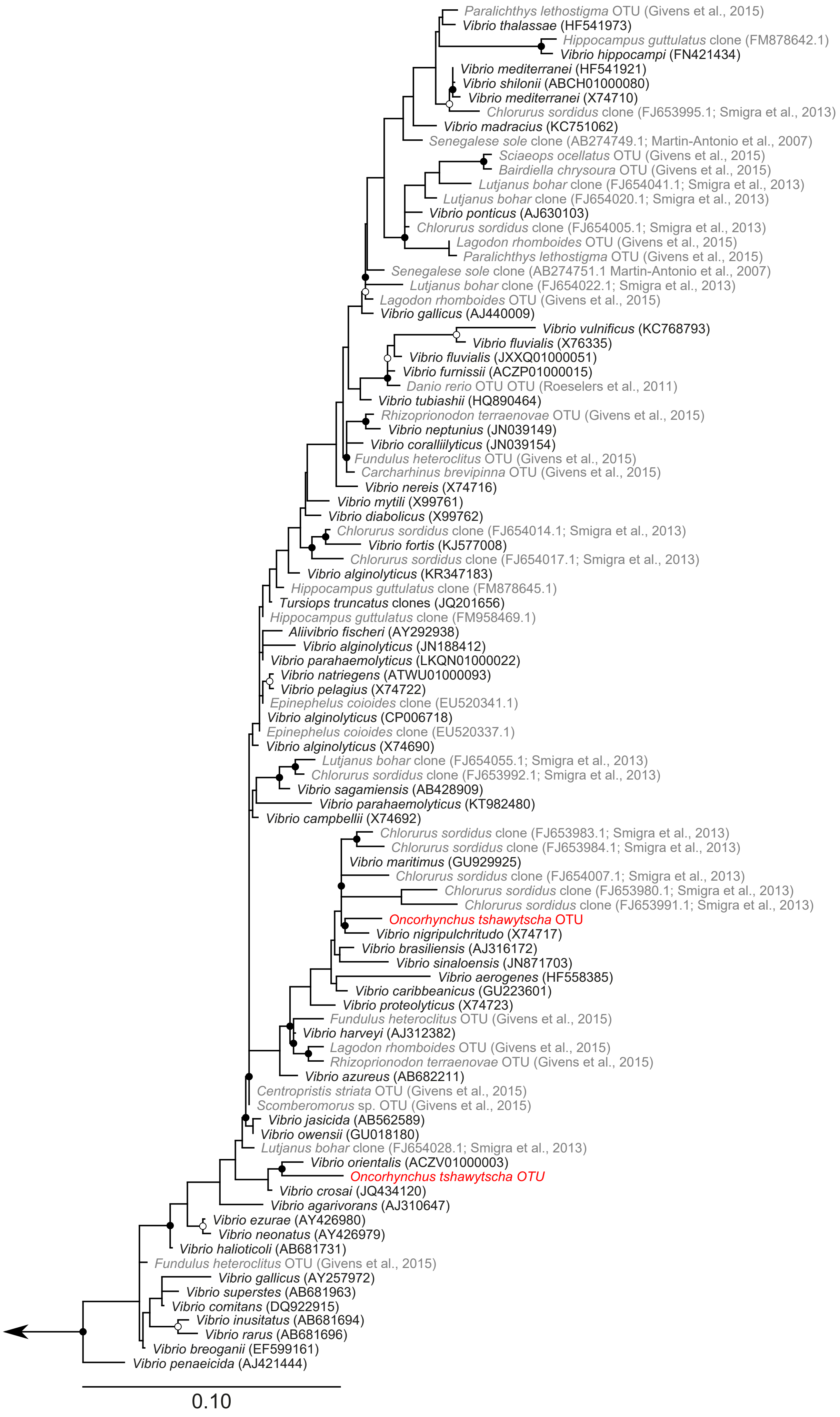

### Supplementary Materials

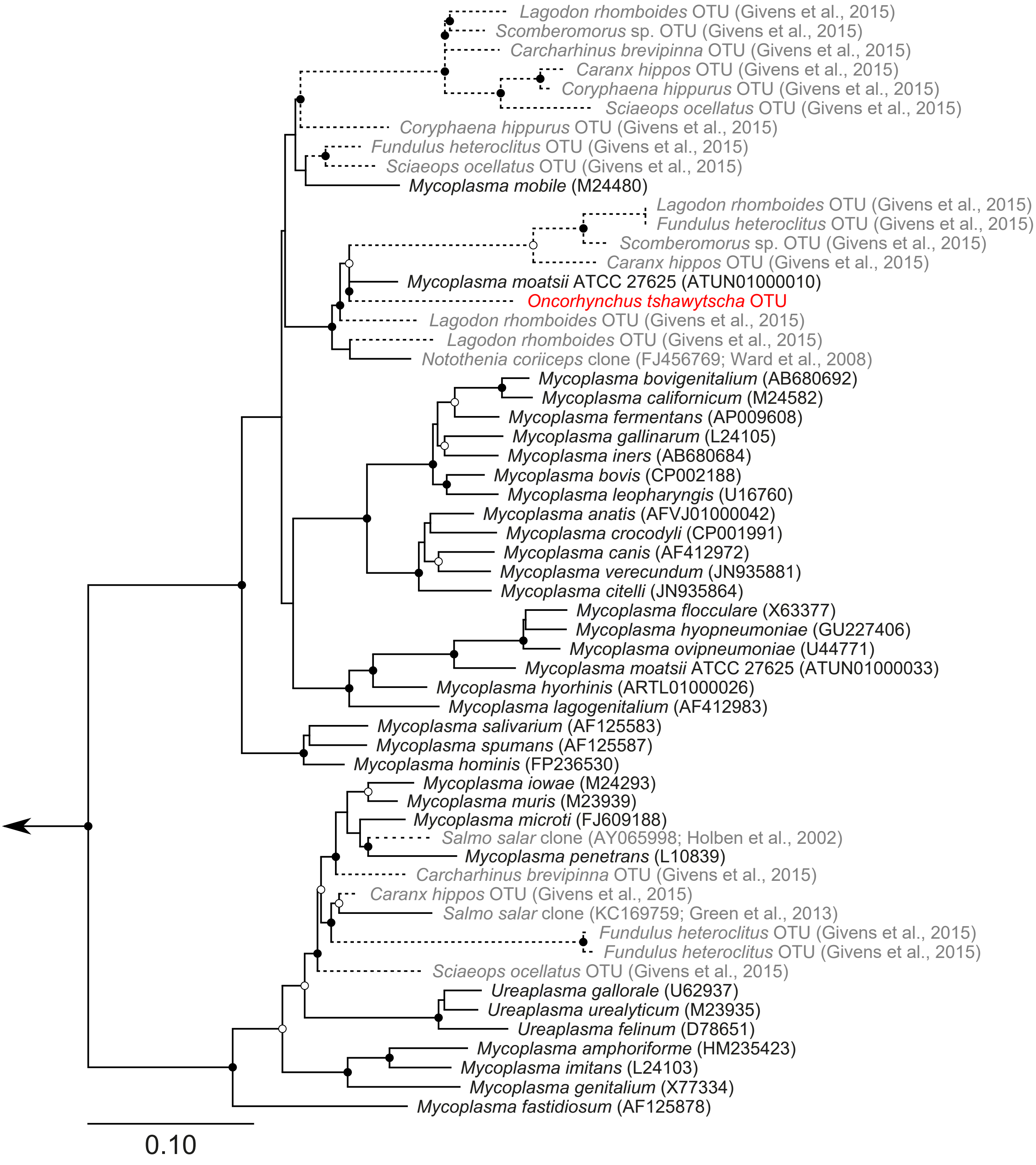

### Supplementary Materials

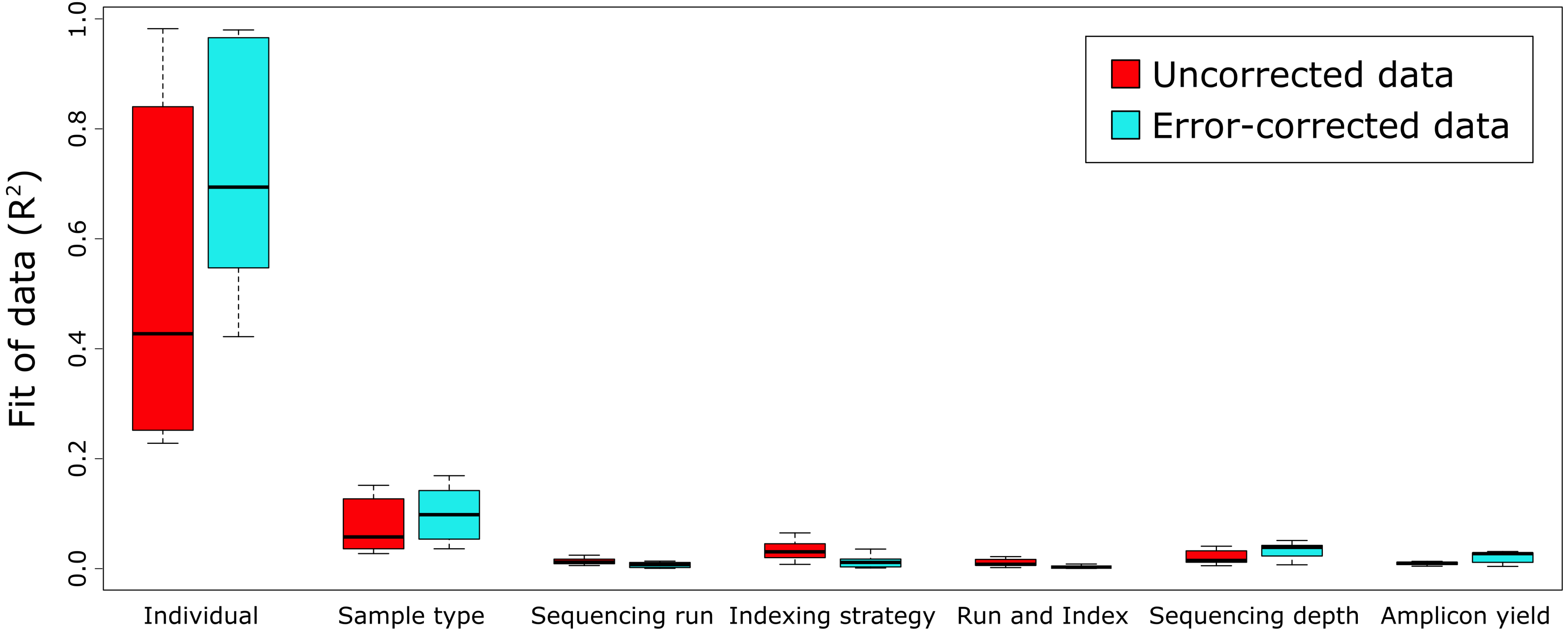

### Supplementary Materials

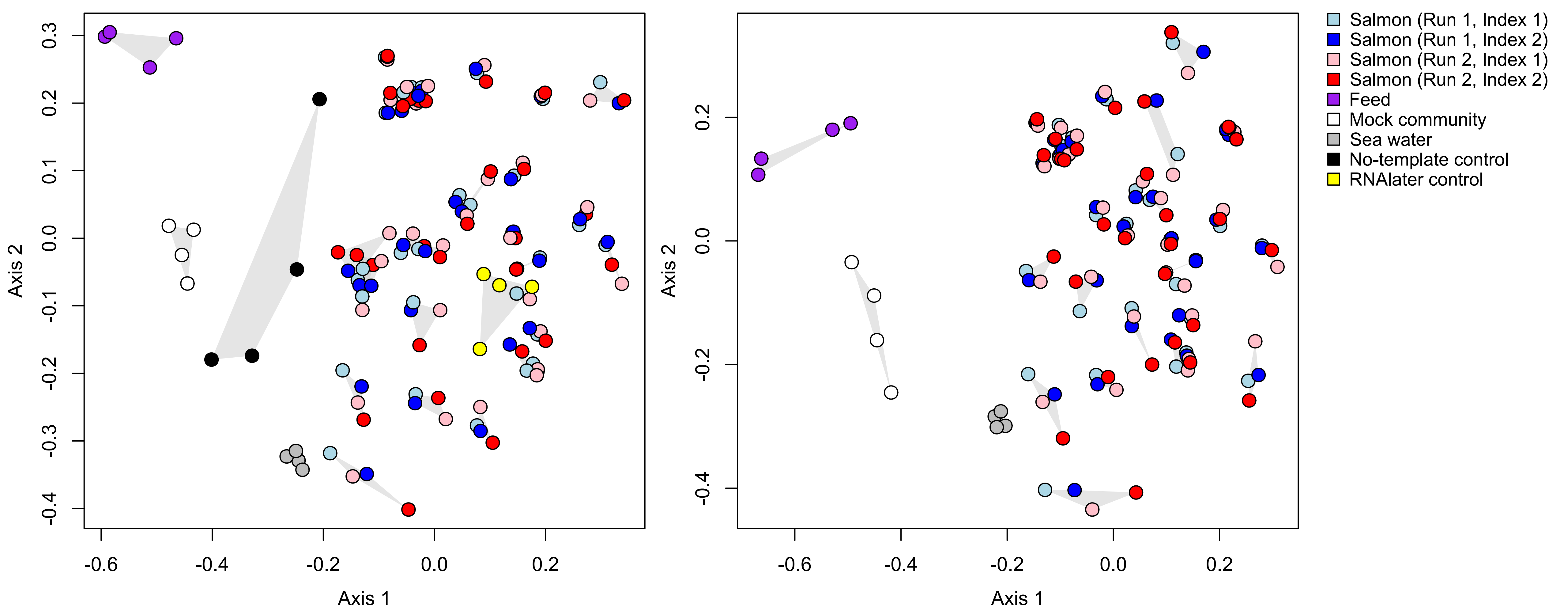

### Supplementary Materials

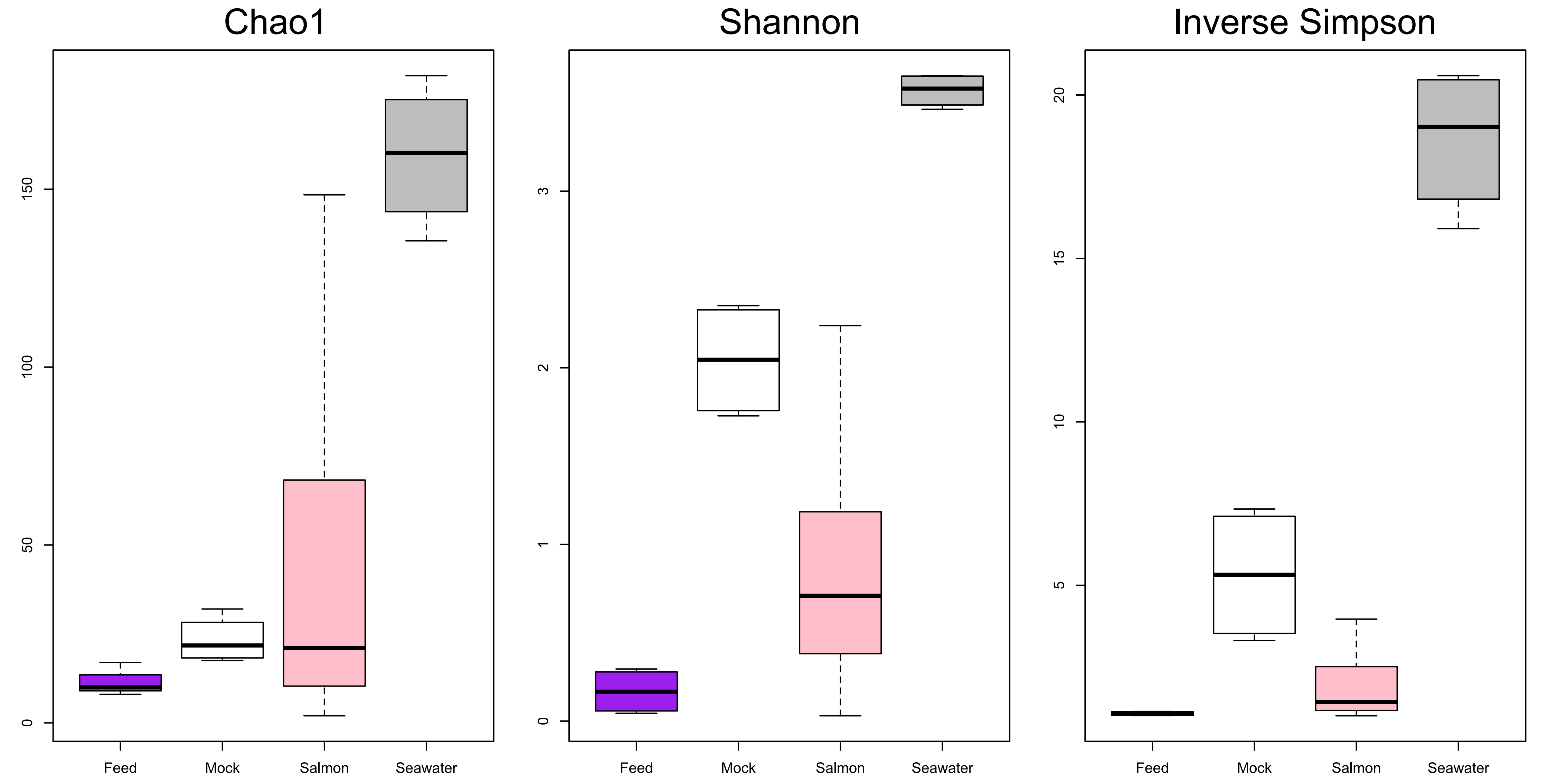

### Supplementary Materials

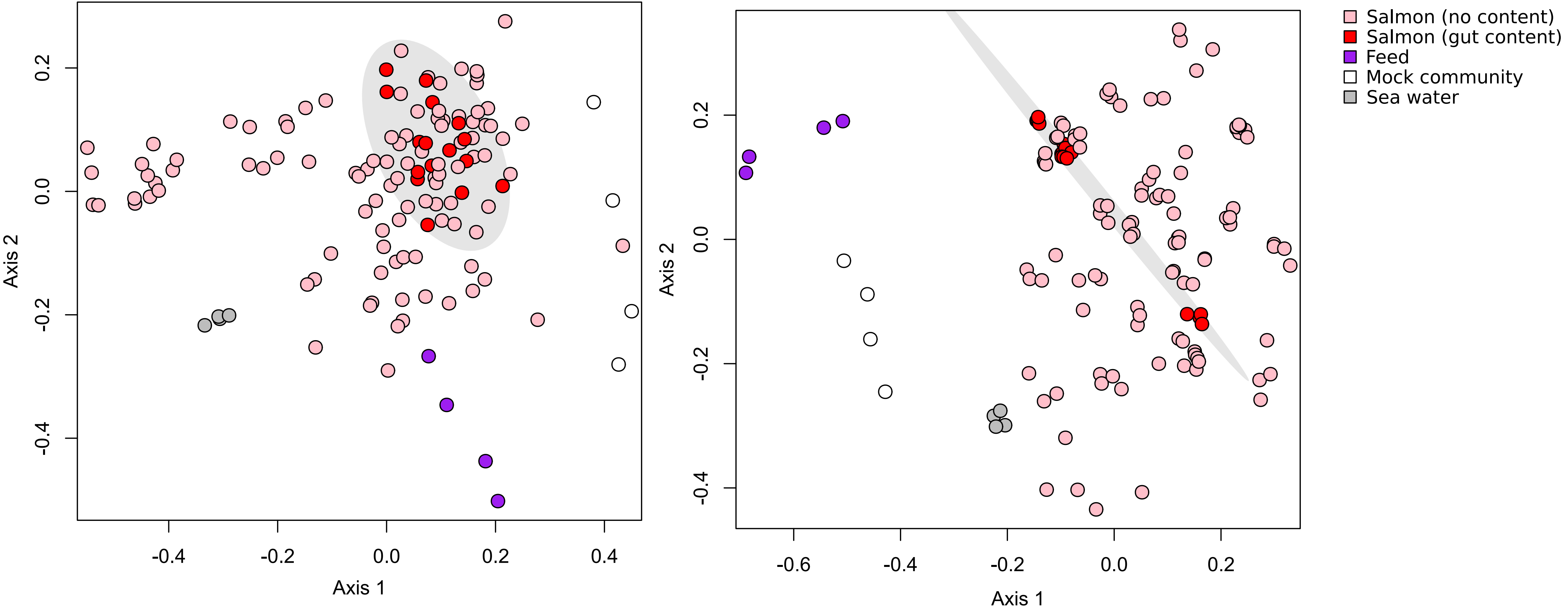
